# CMAPLE: efficient phylogenetic inference in the pandemic era

**DOI:** 10.1101/2024.05.15.594295

**Authors:** Nhan Ly-Trong, Chris Bielow, Nicola De Maio, Bui Quang Minh

## Abstract

We have recently introduced MAPLE (MAximum Parsimonious Likelihood Estimation), a new pandemic-scale phylogenetic inference method exclusively designed for genomic epidemiology. In response to the need for enhancing MAPLE’s performance and scalability, here we present two key components: (1) CMAPLE software, a highly optimized C++ reimplementation of MAPLE with many new features and advancements; and (2) CMAPLE library, a suite of Application Programming Interfaces to facilitate the integration of the CMAPLE algorithm into existing phylogenetic inference packages. Notably, we have successfully integrated CMAPLE into the widely used IQ-TREE 2 software, enabling its rapid adoption in the scientific community. These advancements serve as a vital step towards better preparedness for future pandemics, offering researchers powerful tools for large-scale pathogen genomic analysis.

## Introduction

Phylogenetic analysis plays a vital role in genomic epidemiology, as exemplified during the COVID-19 pandemic (Gonzalez-Reiche et al. 2020; Lu et al. 2020; Hodcroft et al. 2021; Vöhringer et al. 2021). Phylogenetic tools help unveil the origins and transmission of pathogens, monitor the emergence of new variants, and inform vaccine development. For instance, the widely used IQ-TREE 2 software (Minh et al. 2020) has been at the core of the COVID-19 pandemic response, being employed, for example, in Nextstrain (Hadfield et al. 2018). To deal with the pandemic, we recently introduced MAPLE (De Maio et al. 2023), a novel likelihood-based phylogenetic inference method tailored for genomic epidemiological analyses. To address the ever-looming threat of new pandemics, the need for further improvements regarding both the performance and scalability of MAPLE has become increasingly apparent.

Here, we present CMAPLE, a C++ reimplementation of MAPLE highly optimized for performance and scalability. CMAPLE is three-fold faster and more memory efficient than MAPLE, reducing the runtime to analyze 200,000 SARS-CoV-2 sequences (McBroome et al. 2021) using one CPU core from 2.4 days to 19 hours. Notably, CMAPLE can reconstruct a phylogenetic tree of one million SARS-CoV-2 genomes, taking 11 days and 15.4 GB RAM. Additionally, we incorporate a plethora of new protein models (Minh et al. 2021; Dang et al. 2022) to improve CMAPLE’s versatility in analyzing a broader spectrum of pathogen genomic data. We also developed a suite of Application Programming Interfaces (API) and have successfully incorporated CMAPLE into IQ-TREE version 2.3.4.cmaple (https://github.com/iqtree/iqtree2/releases). Additionally, we developed an adaptive mechanism that automatically selects CMAPLE or IQ-TREE search algorithms to minimize the runtime. These advancements facilitate rapid dissemination and widespread adoption of CMAPLE.

To efficiently control pandemics, authorities need to make urgent decisions, such as applying new preventive measures to tackle new virus variants. CMAPLE enables a quicker and more accurate tracking of transmission and discovery of new variants and mutations, which is critical for public health decisions, vaccine designs, and better preparedness for future pandemics.

In the following, we highlight the improvements and key new features in CMAPLE and discuss a few potential directions for further enhancements.

### The overall workflow of the CMAPLE tree search algorithm

CMAPLE is a C++20 reimplementation of the MAPLE algorithm (v0.1.4) implemented in Python. MAPLE takes advantage of low sequence divergence in pathogen data to optimize memory usage. More specifically, MAPLE compresses an input sequence alignment in FASTA or PHYLIP format into the so-called MAPLE format (De Maio et al. 2023), which represents each sequence by a set of the differences to a reference sequence. The reference sequence can be specified by users or automatically computed from the input alignment as the consensus sequence. This results in a very compact data structure, and we have taken a similar approach for the phylogenetic likelihood vectors, allowing much faster (although more approximate) likelihood calculation than the standard Felsenstein’s pruning algorithm (Felsenstein 1973). Our new implementation of CMAPLE focuses on further reducing memory allocation operations, optimizing memory access patterns, and minimizing CPU cache misses.

The CMAPLE algorithm involves three main steps: (1) building an initial tree using the sample placement algorithm (see below), (2) optimizing the tree topology using SPR moves (Felsenstein 1989; Swofford and Olsen 1990), and finally (3) optimizing the branch lengths. Users can choose to skip steps 2 and 3 (`-search fast`) to speed up the analysis, in which case CMAPLE operates like USHER (Turakhia et al. 2021). CMAPLE also supports updating a user-provided input tree, which might not contain all taxa. In this case, the first step of CMAPLE will add the missing sequences to the existing tree using the placement algorithm, then the second step only applies SPR moves to the newly added sequences. Users can choose to skip it (`-search fast`) or to perform a thorough search considering all SPR moves (`-search exhaustive`).

### CMAPLE implementation and performance optimization

In general, high-performance computing and optimization is a very broad topic with multiple factors such as hardware capabilities, programming language, compiler version, choice of algorithm, and data structure, to name a few. We applied many code optimization techniques to enhance the time and memory efficiency of CMAPLE. In brief, while implementing the MAPLE algorithm, we tried to leverage optimization techniques such as SIMD for matrix and vector operations, compact data structures (for tree nodes) with minimal padding and optimal cache-line efficiency (hot/cold splitting), using bitfields and de-virtualization where possible. We also tried to minimize branches, especially in hot loops, to use stack objects instead of heap objects, preallocations and C++ move semantics. It is hard to quantify the individual effect of these techniques since there is interplay between them, yet they all contributed significantly to performance. Last but not least, we employ a high-performance library for memory allocation, jemalloc, to further save runtimes for Linux and macOS. As jemalloc can be activated during runtime, its effect can be quantified more easily and leads to a time saving of up to around 8% (Suppl. Table S2).

CMAPLE relies on three third-party libraries: ncl (https://github.com/mtholder/ncl), simde (https://github.com/simd-everywhere/simde, and zlib (http://zlib.net/).

### Fast (online) sample placement onto an existing tree

During the COVID-19 pandemic, many new viral genome sequences have been obtained and shared on a daily basis. Researchers and healthcare professionals have leveraged fast sample placement onto continually updated and maintained global SARS-CoV-2 phylogenetic trees for identifying new variants, recombinations, and reconstruction transmissions. Therefore, CMAPLE implements a fast sample placement algorithm that allows adding new sequences into a phylogenetic tree of existing samples as follows.

CMAPLE applies the stepwise addition method (Swofford et al. 1996), which iteratively adds new samples to the existing tree one at a time according to their similarity to the reference sequence, such that the most closely related sequence will be added first. For each new sequence, CMAPLE tries to insert it into every branch of the tree and re-evaluates the tree’s likelihood. The branch with the highest likelihood will be chosen. This process is repeated until all new sequences are added to the tree. In our tests, CMAPLE takes 14 minutes to insert 10,000 randomly sampled SARS-COV-2 sequences into an existing 500,000-sample tree. The runtime does not depend on the genome size, but on the divergence level of the added sequences to the reference: a higher divergence level leads to a longer running time.

### Reversible and Non-reversible DNA and Protein Substitution Models

CMAPLE supports two reversible DNA substitution models, JC (Jukes and Cantor 1969) and GTR (Tavaré 1986), and the general nonreversible model UNREST (Yang 1994a). Additionally, we have implemented 40 empirical reversible and non-reversible protein models from the IQ-TREE 2 software (Minh et al. 2021; Dang et al. 2022), which are all listed at https://github.com/iqtree/cmaple/wiki#supported-substitution-models. Those models enable CMAPLE to analyze a broader spectrum of pathogen data, including bacterial genomes (Parks et al. 2018).

### Fast branch tests

Phylogenetic inference typically involves branch support assessment of the inferred trees. To facilitate that task, CMAPLE incorporates the Shimodaira-Hasegawa-like Approximate Likelihood-Ratio Test (SH-aLRT) (Guindon et al. 2010). To take advantage of multi-core CPUs, the SH-aLRT implementation is parallelized using OpenMP (Chapman et al. 2007), which speeds up the SH-aLRT calculation nearly linearly with the number of CPU cores used. For instance, assessing branch support for a phylogenetic tree with 100,000 tips requires 5.1 hours on a single core (AMD EPYC 7551) but only 14 minutes on 32 cores, a 22-fold speedup.

### Application Programming Interface (API) and Integration into the IQ-TREE 2 Software

In addition to a standalone software package, we provide a C++ Application Programming Interface (API) with comprehensive documentation at http://iqtree.org/cmaple/. The CMAPLE API provides three main C++ classes: Alignment, Model, and Tree, representing the input sequence alignment, the substitution models, and the phylogenetic tree, respectively. The API facilitates the integration of the CMAPLE’s algorithm into existing phylogenetic software such as IQ-TREE (Nguyen et al. 2015), RAxML (Stamatakis 2014; Kozlov et al. 2019), and PHYML (Guindon et al. 2010). In fact, we already incorporated CMAPLE into IQ-TREE version 2.3.4.cmaple (https://github.com/iqtree/iqtree2/releases), which users can use via ‘`--pathogen-force`’ option.

### Automatically selecting CMAPLE or IQ-TREE search algorithms to minimize the runtime

The performance of CMAPLE and MAPLE strongly relies on low sequence divergence (De Maio et al. 2023), i.e., the MAPLE algorithm only works well on closely related sequences, such as SARS-CoV-2 genomes. Therefore, we provide a feature to decide if the CMAPLE algorithm is effective for any user-given input alignment: (1) by default, every sequence must be at most 6.7% different from the reference sequence, and (2) the average sequence divergence from the reference must be at most 2%. We set these criteria based on the results of De Maio et al. (2023), who benchmarked MAPLE against popular phylogenetic methods using alignments simulated with different levels of divergence; at levels of divergence around 20 times higher than typical SARS-CoV-2 datasets, they found that MAPLE becomes less accurate or less efficient than traditional approaches, so we set these as thresholds in our method.

Within IQ-TREE version 2.3.4.cmaple, users can use the option ‘`--pathogen`’, which will automatically invoke either the CMAPLE or the original IQ-TREE search algorithm, depending on the effectiveness defined above. For the API, developers can use the function `cmaple::isEffective()`.

### Benchmarking CMAPLE against existing software

We benchmarked the sequential version of CMAPLE (under the default setting and using the jemalloc library) against MAPLE on a server with an AMD EPYC 7551 32-core Processor. To generate a testing dataset, we subsampled 5K, 10K, 50K, 100K, and 200K real SARS-CoV-2 sequences from a global dataset of 4.3 million genomes available on April 2nd, 2022 (McBroome et al. 2021). CMAPLE is about three times faster (Fig. 1A) and requires over three times less memory than MAPLE (Fig. 1B) while yielding equally high likelihood trees as MAPLE (see the “Software Validation” section). This is due to the memory efficiency of C++ code over Python and several optimization techniques (see the “CMAPLE implementation and performance optimization” section). MAPLE required 2.4 days and 11.5 GB RAM (Fig. 1) to reconstruct a tree of 200,000 SARS-CoV-2 sequences, whereas CMAPLE took only 19 hours and 3.6 GB RAM.

**Figure 1.**
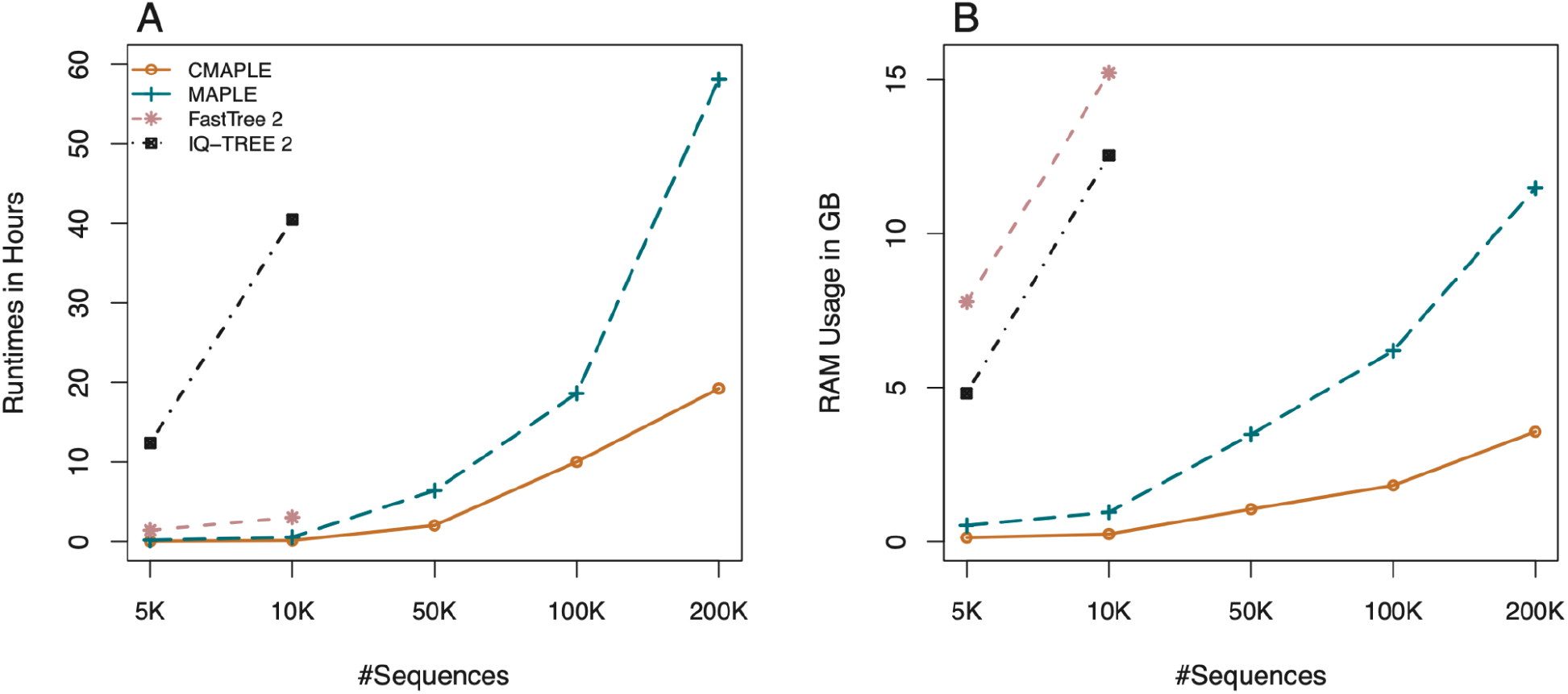
Runtimes (A) and peak memory consumptions (B) of CMAPLE, MAPLE, FastTree 2, and IQ-TREE 2 on analyzing 5K, 10K, 50K, 100K, and 200K SARS-CoV-2 genomes subsampled from a global dataset of 4.3 million genomes available on April 2nd, 2022 (McBroome et al. 2021).

Thanks to these improvements, we ran CMAPLE on an alignment of one million SARS-CoV-2 genomes (McBroome et al. 2021), an input that MAPLE and other existing maximum likelihood methods cannot handle currently. CMAPLE took 11 days and 15.4 GB RAM. Therefore, CMAPLE is highly efficient.

Analyzing these large alignments using existing maximum likelihood software is impractical. Therefore, we benchmarked CMAPLE against IQ-TREE 2 (v2.2.5) and FastTree 2 (Double-precision executable for nearly-identical sequences, v2.1.11) (Price et al. 2010) on two smaller alignments with 5K and 10K real SARS-CoV-2 sequences. CMAPLE is 23x and 300x faster and requires 62 and 51 times less memory than FastTree 2 and IQ-TREE 2, respectively, while also producing trees with higher likelihoods (Suppl. Table S1). CMAPLE took 8 minutes and 0.24 GB RAM to reconstruct a tree from 10K sequences, while FastTree 2 and IQ-TREE 2 took 3 and 40.5 hours and 15.22 and 12.54 GB RAM, respectively (Fig. 1).

### Software Validation

We validated our implementation by using IQ-TREE to compute the likelihoods of the trees inferred by CMAPLE and MAPLE from the real SARS-CoV-2 alignments of our testing datasets above. We found that the likelihoods of the trees inferred by CMAPLE and MAPLE are at most 0.003% different from each other (Suppl. Table S1).

We validated our branch support implementation in CMAPLE with IQ-TREE 2 on the same testing datasets. The SH-aLRT branch supports computed by both programs on the CMAPLE-inferred trees have a Pearson correlation coefficient of 0.995 to 0.997 (Suppl. Fig. S1). We observed 1.98-3.76% and 0.17-0.33% of branches where the two support values differed by more than 10% and 20%, respectively. We anticipate that these differences are due to the approximate nature of the CMAPLE likelihoods.

We also examined the code quality using Softwipe (Zapletal et al. 2021) and achieved an overall absolute score of 8.0/10, ranked third out of 53 computational tools examined at https://github.com/adrianzap/softwipe/wiki/Code-Quality-Benchmark (Access date: 2nd Feb 2024). All datasets and the testing scripts are provided in the Supplemental Material.

### Documentation and User Support

CMAPLE is open-source and freely available at https://github.com/iqtree/cmaple. We provide two executables: “cmaple” and “cmaple-aa” for DNA and protein data, respectively. A comprehensive user manual and command reference of the CMAPLE software are available at https://github.com/iqtree/cmaple/wiki. User support, bug reports, and feature requests can be conveniently submitted at https://github.com/iqtree/cmaple/issues. API documentation and instructions on how to use the CMAPLE library are available at http://iqtree.org/cmaple/.

## Discussions

Existing phylogenetic software, such as IQ-TREE, RAxML, and FastTree 2, was primarily developed for general tree reconstructions, thus, may become slow and inefficient when handling large alignments with closely related sequences, as demonstrated in our benchmark and by De Maio et al. (2023). CMAPLE was specifically designed to address that problem.

We have here presented a sequential tree search algorithm, CMAPLE, which successfully improves upon MAPLE for larger pandemic-scale phylogenetic reconstruction. We plan to further reduce CMAPLE’s runtime by parallelizing tree search using OpenMP (Chapman et al. 2007) and/or message passing interface (MPI) (Gropp et al. 1998). During tree search, CMAPLE seeks SPR moves for internal branches in a sequential manner. To expedite this computationally intensive task, OpenMP can employ multithreading on a single machine, while MPI allows multi-processing in a high-performance computing cluster. Besides, we will also consider other parallelization approaches, such as parallel tree searches, parallelizing the likelihood calculation over genome list entries (i.e., groups of sites, see De Maio et al. (2023)) (similar to the approaches of PLL (Flouri et al. 2015), BEAGLE (Ayres et al. 2012), IQ-TREE, and RAXML), and potentially utilizing GPUs (Smith et al. 2024). These ideas require substantial efforts in design and implementation, thus beyond the scope of the current study.

CMAPLE currently applies GTR for DNA and LG for protein data as default choices. It would be desirable to devise a model selection mechanism to automatically choose the most appropriate substitution model for a given data set, for example, by the Bayesian Information Criterion (Schwarz 1978) and Akaike Information Criterion (Akaike 1974). Besides, the current MAPLE algorithm assumes all sites in the alignment evolve at the same substitution rate. We plan to relax this assumption by implementing models of rate heterogeneity across sites (Yang 1994b).

Another key challenge in genomic epidemiological analysis is to deal with sequencing errors (De Maio et al. 2020; Turakhia et al. 2020). Not accounting for sequencing errors can lead to inaccurate inference (Turakhia et al. 2020). We plan to implement sequence error models (Felsenstein 2004; Chen et al. 2022) to enhance the robustness and accuracy of CMAPLE.

Apart from providing CMAPLE as a standalone program, we also provide it as an API that can be readily deployed in any C++ code. The API provides the `cmaple::isEffective()` function to quickly test the efficiency of the CMAPLE algorithm for a dataset at hand. If efficient, developers can invoke the CMAPLE library as provided in the tutorial. Besides, we plan to create a Python package for CMAPLE to approach a wider user base (e.g., similar to Wang et al. 2023).

The CMAPLE library complements existing phylogenetic libraries, PLL and BEAGLE, but cannot replace them because the CMAPLE algorithm only works on low divergence sequences. In the future, we plan to combine the methods in CMAPLE with classical phylogenetic approaches (such as those implemented within the libraries PLL and BEAGLE) to develop a unifying approach that would work efficiently at any level of divergence, for example, switching from classical algorithms and data structures to those of CMAPLE as one moves from long branches in the tree into densely sampled clades.

## Supplementary Material

The data underlying this article are available in the Zenodo Repository at https://doi.org/10.5281/zenodo.11180100.

## Acknowledgments

This work was supported by a Chan-Zuckerberg Initiative grant for open source software for science to B.Q.M. [EOSS4-0000000312]; an Australian Research Council Discovery grant [DP200103151 to B.Q.M.]; a Moore-Simons Foundation grant [735923LPI (https://doi.org/10.46714/735923LPI) to B.Q.M.]; and partly by a Vingroup Science and Technology Scholarship [VGRS20042M to N.L.T.]. The computational results have been obtained on the cluster at the Center for Integrative Bioinformatics Vienna (CIBIV). We thank Arndt von Haeseler for providing access to the CIBIV cluster; Prabhav Kalaghatgi, Robert Lanfear, and Thomas Wong for their valuable comments and discussions.

