## Supplementary for "CMAPLE: efficient phylogenetic inference in the pandemic era"

**Table S1.** Runtimes, peak memory usage, and log-likelihoods of the trees inferred by four tools, CMAPLE, MAPLE, FastTree 2, and IQ-TREE 2, testing on 5K, 10K, 50K, 100K, and 200K SARS-CoV-2 sequences. IQ-TREE 2 and FastTree 2 were not run on 50K, 100K and 200K due to excessive runtimes. Each test was run three times, and the average results were reported. The best values among the four methods are in **boldface**.

| #sequences | Software | Log-Likelihood | Difference from the best | Runtimes (hours) | Ratio to the shortest runtime | Peak memory usage (GB) | Ratio to the lowest peak memory usage |
|---|---|---|---|---|---|---|---|
| **5K** | CMAPLE | **-240,820** | 0 | **0.049** | 1 | **0.129** | 1 |
| | MAPLE | **-240,820** | 0 | 0.183 | 3.735 | 0.525 | 4.07 |
| | FastTree 2 | -241,748 | -927 | 1.436 | 29.306 | 7.787 | 60.364 |
| | IQ-TREE 2 | -241,739 | -919 | 12.398 | 253.02 | 4.813 | 37.31 |
| **10K** | CMAPLE | **-391,549** | 0 | **0.135** | 1 | **0.244** | 1 |
| | MAPLE | -391,590 | -41 | 0.563 | 4.17 | 0.956 | 3.918 |
| | FastTree 2 | -393,393 | -1,844 | 3.049 | 22.585 | 15.22 | 62.377 |
| | IQ-TREE 2 | -393,492 | -1,944 | 40.488 | 299.911 | 12.542 | 51.402 |
| **50K** | CMAPLE | **-1,274,343** | 0 | **2.036** | 1 | **1.056** | 1 |
| | MAPLE | -1,274,367 | -24 | 6.412 | 3.149 | 3.479 | 3.295 |
| **100K** | CMAPLE | -2,010,053 | -55 | **10.021** | 1 | **1.825** | 1 |
| | MAPLE | **-2,009,998** | 0 | 18.631 | 1.859 | 6.199 | 3.397 |
| **200K** | CMAPLE | **-3,425,616** | 0 | **19.236** | 1 | **3.569** | 1 |
| | MAPLE | -3,425,673 | -57 | 58.094 | 3.020 | 11.476 | 3.215 |

**Table S2.** Runtimes, peak memory usage, and log-likelihoods of the trees inferred by CMAPLE executed with the high-performance memory allocator - jemalloc (CMAPLE) and without jemalloc (CMAPLE_wo_jemalloc). The tests were conducted on 5K, 10K, 50K, 100K, and 200K SARS-CoV-2 sequences. Each test was run three times, and the average results were reported.

| #sequences | Versions | Log-Likelihood | Difference from the best | Runtimes (hours) | Time saving compared with CMAPLE_ wo_jemalloc | Peak memory usage (GB) | Ratio to the lowest peak memory usage |
|---|---|---|---|---|---|---|---|
| 5K | CMAPLE | -240820.106 | 0.000 | 0.049 | 0.00% | 0.129 | 1.000 |
| | CMAPLE_ wo_jemalloc | -240820.106 | 0.000 | 0.049 | | 0.129 | 1.000 |
| 10K | CMAPLE | -391548.681 | -0.009 | 0.135 | 0.20% | 0.244 | 1.000 |
| | CMAPLE_ wo_jemalloc | -391548.672 | 0.000 | 0.136 | | 0.244 | 1.000 |
| 50K | CMAPLE | -1274343.252 | -0.003 | 2.036 | 0.22% | 1.056 | 1.000 |
| | CMAPLE_ wo_jemalloc | -1274343.249 | 0.000 | 2.040 | | 1.056 | 1.000 |
| 100K | CMAPLE | -2010053.299 | 0.000 | 10.021 | 7.71% | 1.825 | 1.000 |
| | CMAPLE_ wo_jemalloc | -2010053.302 | -0.003 | 10.859 | | 1.824 | 1.000 |
| 200K | CMAPLE | -3425615.639 | 0.000 | 19.236 | 4.72% | 3.569 | 1.001 |
| | CMAPLE_ wo_jemalloc | -3425616.251 | -0.612 | 20.189 | | 3.567 | 1.000 |

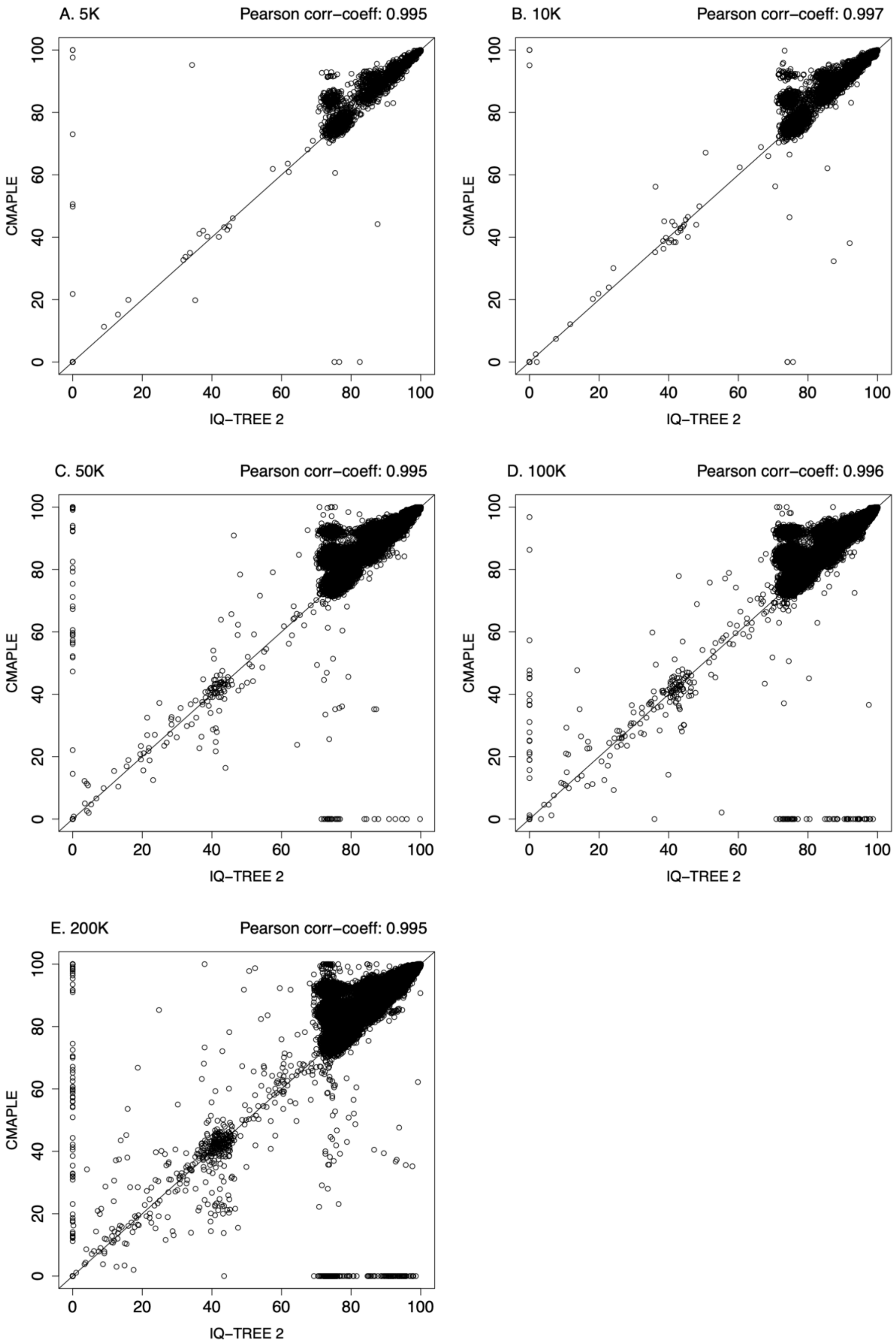


**Figure S1.** Branch supports computed by CMAPLE and IQ-TREE 2 of trees inferred by CMAPLE from 5K (sub-panel A), 10K (sub-panel B), 50K (sub-panel C), 100K (sub-panel D), and 200K (sub-panel E) real SARS-CoV-2 sequences.
